# Social Distance and Fungal Disease in an Australian Wild Lizard Population

**DOI:** 10.64898/2026.04.06.716608

**Authors:** Félix Requena-García, Nicola Jackson, Barbara Class, Angela C. Mitchell, Rebecca L. Cramp, Celine H. Frere

## Abstract

Social living often confers substantial fitness benefits; however, close spatial association among individuals can also elevate opportunities for pathogen transmission, especially where the populations are dense. Despite this, the extent to which avoidance behaviours are expressed by wild reptiles facing fungal disease remains unclear. We examined Eastern Water Dragons (*Intellagama lesueurii*) in Roma Street Parklands, Brisbane, Australia, where a population is affected by the emerging fungal pathogen *Nannizziopsis barbatae* (*Nb*). Using a five-year dataset (2018-2023) spanning 146 individuals, we quantified social distance, as the minimum distance to the nearest neighbour, in relation to the number of diseased conspecifics that overlapped each individual’s seasonal core home area. Social distance was negatively associated with the number of diseased conspecifics, indicating a strong crowding effect; however, this association was weaker for diseased individuals, which were observed at larger distances than healthy individuals even when the number of diseased conspecifics was high. Together, these patterns indicate a context-dependent spacing response associated with disease status and local disease context, suggesting that infection risk constrains density-driven proximity. Our findings provide new insights into how disease pressure shapes social spacing in reptiles and contribute to a broader understanding of behavioural responses to emerging infectious fungal diseases.

## Introduction

In behavioural ecology, sociality is thought to evolve when the benefits associated with group living exceed its costs (Silk, 2007). Numerous studies have shown that sociality can confer important fitness advantages, such as reducing predation risk and increasing mating opportunities (Alexander, 1974; Silk, 2007; Van Schaik, 1983). However, sociality can simultaneously increase infection risk (Dobson, 1988; Manlove et al., 2014), as the proximity and frequent interactions among group members facilitate pathogen transmission (Romano et al., 2020). Consequently, how animals respond to diseased conspecifics is critical for individual survival and can shape pathogen transmission dynamics at the population level (Townsend et al., 2020).

Infectious diseases can have profound effects on population dynamics and species persistence (Altizer et al., 2003; Curtis, 2014; Fong, 2017; Hart and Hart, 2021; Kappeler et al., 2015; Langwig et al., 2012; Maher et al., 2012). Emerging infectious fungal diseases, in particular, have had far-reaching impacts on both wildlife and humans, with millions of infections and deaths recorded annually (Denning, 2024; Fisher et al., 2012). A well-known case is chytridiomycosis in amphibians associated with the fungus *Batrachochytrium dendrobatidis*, driving large-scale population declines and the erosion of biodiversity worldwide (Scheele et al., 2019). Compared with many parasites and bacterial or viral pathogens, fungal pathogens present unique conservation challenges because they can persist in the environment independently of hosts and spread through both environmental reservoirs and direct host-to-host contact, reducing the dependence of transmission on host density alone (Bahram and Netherway, 2022; Fisher et al., 2012).

When infection risk is high, animals that live in groups may alter their behaviour to reduce exposure, with avoidance strategies playing a key contribution to limiting transmission (Croy et al., 2023; Hoekendijk et al., 2023). Such strategies include social distancing, where healthy (and occasionally sick) individuals avoid close contact with conspecifics showing signs of illness, and self-isolation, in which individuals reduce interactions to conserve energy or limit competition (Bos et al., 2012; Rueppell et al., 2010). Avoidance behaviours have been documented across multiple taxonomic groups (Lopes et al., 2022), while self-isolation has been most clearly demonstrated in eusocial insects, and, to a lesser extent, in vertebrates such as badgers and immunocompromised mice (Lopes et al., 2022). However, in wildlife systems dominated by fungal pathogens, where both environmental and direct transmission pathways may operate simultaneously, the role of behavioural avoidance remains poorly understood. In such systems, spatial and social behaviours are tightly coupled, making it difficult to disentangle their relative constraints on transmission dynamics (Albery et al., 2024). This leads to the question of how effective avoidance behaviour can be in reducing infection risk when environmentally mediated transmission constrains the possibility of complete avoidance.

Navigating a diseased social landscape requires individuals to make decisions not only about whom to avoid but also about whom to continue socialising with (Weinstein et al., 2018). Although earlier studies have largely focused on the avoidance of diseased individuals, it remains unclear whether such responses are expressed consistently or only under specific conditions, a phenomenon often described as partial avoidance. Decisions to avoid infected conspecifics may depend on infection severity, duration of exposure risk (Croft et al., 2011; Tacey et al., 2023), sex (Paciencia et al., 2019), pathogen type (Manlove et al., 2014; Paciencia et al., 2019), or chemical cues (Kiesecker et al., 1999). We refer to this behavioural contingency as context-dependent avoidance, whereby spacing responses vary according to local disease information rather than being expressed uniformly. Here partial avoidance refers to decreases in proximity or contact frequency that occur without full cessation of social interaction, representing a potential mechanism through which individuals balance the costs and benefits of sociality under elevated disease risk. Previous work on eastern water dragons has shown that fungal pathogens can be transmitted via environmental reservoirs, suggesting that social avoidance alone may be insufficient to fully prevent infection (Becker et al., 2025; Tacey et al., 2023). Nevertheless, behaviour may still play a key role in shaping exposure risk. This raises the question of how individuals adjust their spatial and social behaviour when complete avoidance is constrained by environmentally mediated transmission.

Based on this framework, we hypothesise that social distancing, specifically how adjustments in social distance in response to infection risk, represents a key form of partial avoidance in diseased populations. Rather than eliminating exposure, such behavioural responses may modulate infection risk under conditions where environmental transmission limits complete avoidance. We predict that social distancing will be influenced by the number of diseased conspecifics (Tacey et al., 2023), and that increasing crowding will constrain opportunities for maintaining distance, as increased crowding typically promotes more frequent interactions (Strickland and Frere, 2019). Conversely, we expect social distance to increase under heightened disease risk, reflecting avoidance responses triggered by the presence of more diseased conspecifics.

We tested these hypotheses using five years of data from a long-term (>14 years) research project on an eastern water dragon population in Roma Street Parklands, a system well-suited to the study of social plasticity due to its high sociality and sensitivity to environmental change (Peterson et al., 2021; Piza-Roca et al., 2019; Piza-Roca et al., 2018; Strickland and Frere, 2019; Strickland et al., 2021). In 2013, an outbreak of *Nb*, a fatal fungal pathogen previously identified only in captive reptiles, emerged at Roma Parklands and now infects a substantial proportion of the population (Becker et al., 2025; Peterson et al., 2021). Disease prevalence has increased over the past decade, rising from relatively low levels following emergence to affecting more than half of the monitored population (21.1% in 2018 to 62.4% in 2023). Previous work has linked *Nb* infection with elevated mortality risk (Class et al. 2021, unpublished bioRxiv preprint). Building on previous findings based on short-term exposure and dyadic interactions (Tacey et al., 2023), our study leverages a longitudinal dataset (2018-2023) of 146 individuals to examine how social spacing varies across seasons and disease contexts. By focusing on social distance, defined as the minimum distance to the nearest neighbour, we provide a population-level perspective on disease-related social behaviour beyond fine-scale dyadic approaches.

## Methods

### Study population

The parklands population of dragons in Brisbane (27°27’46’S, 153°1’11’E), Australia (Gardiner et al., 2014) comprises more than 300 individuals (Strickland et al., 2014). The 16-ha park features several microhabitats that vary in size, complexity, vegetation structure and proximity to water sources (Littleford-Colquhoun et al., 2019). Moreover, the park is enclosed by roads and other forms of infrastructure, effectively restricting movement between adjacent populations (Littleford-Colquhoun et al., 2017). Eastern water dragons exhibit site fidelity, overlapping home ranges, and agonistic interactions, particularly among males (Baird et al., 2012; Strickland et al., 2014). The behavioural dataset spans September to April of 2018-2023, corresponding to the active season and the years when *Nb* was diagnosed and quantified by qPCR (Peterson et al., 2021; Powell et al., 2021).

### Ethical clearance

All procedures involving animals complied with institutional ethical guidelines and received approval from the Animal Ethics Committee of the University of Queensland (2022/AE000453), and the University of the Sunshine Coast (ANS1858 and ANA20161). Along with permits from Queensland Parks and Wildlife Service (QPWS), research in the area was conducted under authorisation granted by Partnerships permits (P-PTUKI-100129158).

### Behavioural data collection

Data collection was carried out through a long-running behavioural study of Roma Parklands eastern water dragon population initiated in 2010. From September to April of each year, visual surveys were conducted approximately three times per week during morning (7.30-10.30) and afternoon (13.00-16.00) sessions, covering ∼ 85% of the identified population (Littleford-Colquhoun et al., 2019; Piza-Roca et al., 2019; Piza-Roca et al., 2018; Strickland et al., 2021). Individual identification was performed using head profile photographs taken with a Canon EOS 600, based on facial patterning (Gardiner et al., 2014). Spatial locations were obtained using a handheld GARMIN eTrex10 GPS unit. Behaviours, such as resting (inactive, often shaded), basking (inactive, sun-exposed), and agonistic interactions (e.g., fighting, head bobbing, arm waving) were recorded (Baird et al., 2012). This population has been continuously monitored since 2010, and individuals are habituated to both human presence and research activities. As a result, animals typically do not alter their behaviour in response to slowly approaching observers, reducing the likelihood that recorded spatial relationships were influenced by observed disturbance. This approach helps ensure that measured distances reflect natural social interactions rather than artefacts of data collection.

Sex was classified using sexual dichromatism together with secondary sexual characteristics of this species (Strickland et al., 2014). To distinguish adults from juveniles, we followed Tacey et al. (2023). Specifically, this classification relies on several morphological features, including snout-vent length (females > 171 mm, males > 205 mm), eye-ear distance, eye size, and evidence of scarring or missing head crest spikes. In comparison to juveniles, adults exhibit longer, sharper spikes and more pronounced colouration (Tacey et al., 2023). Only individuals with available SVL measurements were retained in the dataset. By applying this approach, we ensured that only adult individuals were included in our study, given that juveniles exhibit distinct social behaviour (Delmé et al., 2024). Individuals were identified with I3S Manta from profile photos based on scale/colour patterns, followed by manual validation (Gardiner et al., 2014; Van Tienhoven et al., 2007).

### Disease status assessment

Biannual dragon captures were conducted throughout the study period (2018 – 2023) to monitor disease. The dragons were captured using a lassoing method as outlined in previous work (Littleford-Colquhoun et al., 2017). Disease status in Roma Parklands dragons was assessed according to the presence of yellowish-brown crusted lesions on the skin, indicative of *Nb* infection (Figure S1). To further validate the presence of the pathogen (*Nb*) in both lesioned and asymptomatic animals, skin swabs were collected by gently rubbing the skin surface with a saline-moistened sterile swab for up to 30 s. All samples were stored at -20 °C before undergoing diagnostic qPCR analyses for *Nb*.

DNA extraction from skin swabs was performed using DNeasy Plant Pro kits (Qiagen). Presence/absence of *Nb* was assessed using a species-specific qPCR assay (Powell et al., 2023). The following primers were used: forward 5’-TGATCATGTTTAGTCTCTGAAGGT-3’ and reverse 5’-AGGCTAAGCTGATTTGGTCTC-3’. qPCR reactions consisted of 10 µl (2X) GoTaq Enviro (Promega), 0.4 µM each of forward primer and reverse primer (10nM concentration), 0.1 µM of the probe 5’-6-FAM/TCCCTGCTG/ZEN/ATTGCCATATATTAGGT/FQ/-3’, and 2 µl template DNA, made up to a total reaction volume of 20 µl using molecular grade water. All qPCR reactions were run in duplicate, and positive controls and no-template controls (DNA/RNA free water) were included in all runs. Samples were considered positive when amplification crossed the cycle threshold (Ct) in both replicates, as determined by CFX96 manager software (Bio-Rad Laboratories, Hercules, CA, USA), under the following thermocycling conditions: 95 °C for 5 min, followed by 40 cycles of 95 °C for 20 s, and 63 °C for 30 s.

The behavioural dataset was restricted to animals for which disease status could be definitively determined, using a combination of visual assessment during capture and molecular diagnostics. Individuals were classified as diseased when they exhibited visual signs of *Nb* infection (e.g., discoloured scales and/or skin lesions) and tested qPCR-positive. Individuals were classified as healthy when they showed no visual signs of *Nb* infection and were qPCR-negative (Tacey et al., 2023). Asymptomatic but qPCR-positive individuals (i.e., individuals with confirmed infection but no visible clinical signs) were identified in the dataset. These individuals may represent early-stage infections or low pathogen loads that have not yet resulted in visible disease (Grabow et al., 2024), although their classification remains inherently uncertain. While asymptomatic individuals represented a relatively small proportion of observations (10%), we included them in an additional model as a separate category to explore potential differences in spacing behaviour. This analysis is described in the Results and reported in Table S7.

### Quantifying core home ranges and conspecific count

GPS data collected during the behavioural surveys were used to estimate per-individual-season core home ranges within kernel utilisation distribution (UDs) implemented in the *adehabitatHR* package (Calenge, 2006). A fixed smoothing parameter of 7 m was applied (Gardiner et al., 2014). Core home ranges were defined using the 50% utilisation distribution contour, representing the area where individuals spend most of their time and where the most social interactions take place (Piza-Roca et al., 2018; Strickland et al., 2017). To ensure stable home-range estimation, a minimum of 30 sightings per individual per season was required (Strickland et al., 2017). Although estimates stabilise with around 20 sightings, we used a stricter threshold of 30 sightings to maximise confidence. As a robustness check, we re-ran the model using a lower threshold of 25 sightings (Table S4). Following data filtering, including restriction to adult individuals with known disease status and at least 30 sightings per season for reliable home-range estimation, the final dataset used in the primary analysis comprised 146 individuals and 10,583 social-distance observations.

For each focal individual and season, we counted unique conspecifics whose GPS locations fell within the focal individual’s 50% core home range. From these counts, we derived *DiseasedCount*, defined as the subset of conspecifics classified as diseased during that season. Core home ranges and conspecific counts were calculated separately for each season to capture temporal variation and avoid cross-seasonal overlap. All analyses were conducted in R (version R.4.2.2; R Core Team (2024). Total conspecific count was excluded because it was highly correlated with *DiseasedCount* (r = 0.88), creating substantial collinearity between predictors. Because diseased individuals tend to occur in high-use areas, *DiseasedCount* is likely to capture aspects of both local crowding and disease exposure. Consequently, the effects estimated should be interpreted as reflecting broader disease-associated social context rather than disease exposure alone.

### Response variable - social distance

Social distance was quantified as the minimum distance (m) to the nearest neighbour per survey session (morning and afternoon). For each survey, a single social-distance measure was obtained for each focal individual from GPS positions recorded during the same survey session. Smaller values indicate higher social tolerance or gregariousness, whereas larger values indicate lower tolerance. All behavioural observations of focal individuals were included, with no minimum sighting threshold per survey (Class et al., 2024). Unlike binary social metrics (e.g., “gregarious” vs “non-gregarious”) (Strickland and Frere, 2019; Strickland et al., 2014), this continuous measure captures a spectrum of social spacing and provides higher resolution and statistical power. Social distance was strongly negatively correlated with the binary gregariousness index described by Class et al. (2024).

### Censoring model structure

Social distance represents the minimum observable distance to a nearest neighbour within a survey and is therefore subject to left censoring due to the GPS resolution limit. Spatial positions were recorded using handheld GPS devices, which introduce a degree of location uncertainty, particularly when individuals are recorded in close proximity. As a result, very small observed distances may reflect measurement imprecision rather than true physical contact. Distances smaller than 1.64 m cannot be reliably distinguished and are recorded at this minimum observable value. To correctly account for this measurement constraint, we modelled these observations using a left-censored likelihood following Class et al. (2024). In this framework, distances below the detection threshold are treated as censored rather than as exact values, which propagates uncertainty about true sub-threshold distances and avoids biasing estimates towards artificially small values. Censoring was implemented directly in the likelihood using the *cens()* argument in the *brms* package (Bürkner, 2017, 2021).

### Statistical analysis

We analysed social distances using a Gamma-distributed generalised linear mixed model fitted with a log link and left-censoring, to account for the strictly positive, right-skewed distribution of distances and the lower detection limit imposed by GPS precision (Class et al., 2024). Because the Gamma distribution cannot accept zero or threshold-level values, we assigned observations at or below 1.64 m a small positive constant before model fitting, while still modelling them explicitly as left-censored. The size of this constant was evaluated via simulation under an intercept-only Gamma model, and a threshold-equivalent value (1.0 × detection limit) yielded the lowest Root mean square error (RMSE) across replicates (Table S1).

Fixed effects included focal diseased status, sex, number of sightings, log-transformed core home range area (m^2^), the number of diseased conspecifics present within the focal individual’s core home range (*DiseasedCount*), and the interaction between focal disease status and *DiseasedCount*. Repeated measurements were accommodated by including random intercepts for individual identity (Name) and individual-season (Name/Season) grouping. The Season component was included to account for unexplained variation among field seasons rather than to estimate temporal trends directly. These random effects account for repeated measures of individuals across seasons and control for individual-level heterogeneity. All continuous count predictors were z-transformed before analysis to improve comparability of effect sizes and reduce collinearity. Models were fitted using default, weakly uninformative priors. The primary model was run with four Markov chains of 4,000 iterations each, including 2,000 warm-up iterations; supplementary sensitivity models used additional iterations where required. Model diagnostics are provided in the supplementary material, including collinearity checks (Table S2), Stan trace plots (Figure S2), posterior predictive checks for censoring behaviour (Figure S3), and PSIS-LOO cross-validation diagnostics (Figure S4).

To assess whether the observed interaction could reflect general density or the presence of non-diseased individuals, we fitted additional models that replaced *DiseasedCount* with the total conspecific count (*TotalCount*) and the healthy conspecific count (*HealthyCount*). We also fitted an exploratory model distinguishing asymptomatic and symptomatic individuals. These analyses were conducted using the same model structure and sampling settings as the main analysis (Tables S5-S7).

To assess the robustness of home-range estimation and model inferences to the minimum sighting threshold, we performed a sensitivity analysis using a reduced inclusion criterion of ≥25 sightings per individual per season. The model structure was identical to the primary analysis (≥30 sightings). Full results are presented in Table S4.

### Model performance

Model adequacy was assessed using posterior predictive checks tailored to the censored Gamma model structure. The model accurately reproduced the observed distribution of social distances, including the long right tail. Convergence diagnostics indicated good performance with R-hat ≤ 1.01 and bulk and tail effective sample sizes > 1,000. Multicollinearity among predictors was low (all VIF < 2.39). For the supplementary sensitivity analyses, additional post-warm-up iterations were used as needed to ensure that the bulk and tail effective sample sizes also exceeded 1,000.

## Results

### Summary statistics

The dataset comprised 522 behavioural surveys yielding 10,583 behavioural records from 146 unique individuals. Summary statistics describing sampling effort, population composition, disease status, observation effort, social-distance measures, diseased conspecific abundance and home-range characteristics are provided in Table 1.

**Table 1:** Summary statistics describing the study population, sampling effort, disease status, observation effort, social-distance measures, diseased conspecific abundance, and 50% core home-range characteristics of Eastern Water Dragons included in the analysis. Values are presented as means ± SD unless otherwise indicated. Diseased conspecific abundance is reported as median and interquartile range (IQR)

| Variable | Value |
| --- | --- |
| Behavioural surveys | 522 |
| Behavioural records | 10,583 |
| Unique individuals | 146 |
| Males | 65 |
| Females | 81 |
| Healthy individuals | 69 (47.3%) |
| Diseased individuals | 77 (52.7%) |
| Observations per individual-season | 52.55 ± 17.55 (range: 30-102) |
| Individuals observed across multiple field seasons | 30.8% |
| 50% core home-range area (m<sup>2</sup>), males | 611 ± 270 |
| 50% core home-range area (m<sup>2</sup>), females | 366 ± 112 |
| Social distance (m), males | 9.97 ± 9.3 |
| Social distance (m), females | 8.19 ± 8.05 |
| Diseased conspecific count, median (IQR) | 8 (IQR: 4-16) |

### Effects on social distance

Posterior estimates showed clear associations of both focal disease status and disease context with social distance (Table 2). Nearest-neighbour distance was greater to diseased focal individuals than to healthy focal individuals (posterior mean = 0.11, 95% CrI [0.03 to 0.20]), consistent with greater social spacing among diseased individuals. Regardless of focal disease status, diseased conspecific count was negatively associated with social distance (posterior mean = -0.30, 95% CrI [-0.39 to - 0.21]), with individuals observed at shorter distances in areas characterised by a higher number of diseased conspecifics.

**Table 2.** Posterior summaries from the Bayesian censored-Gamma mixed-effects model examining social distance (nearest-neighbour distance, m) in eastern water dragons in relation to focal disease status, the number of diseased conspecifics, their interaction, sex, sighting frequency, and core home-range area. Parameter estimates are presented on the log scale because a log-link function was used in the Gamma model.

| Parameter | Posterior Mean | Posterior (SD) | 95% CrI | R-hat |
| --- | --- | --- | --- | --- |
| **Intercept** | **2.021** | **0.052** | **[1.918, 2.123]** | **1.000** |
| Sex (Male) | 0.032 | 0.076 | [-0.115, 0.182] | 1.000 |
| **Diseased (Yes)** | **0.115** | **0.042** | **[0.031, 0.195]** | **1.000** |
| **DiseasedCount** | **-0.303** | **0.045** | **[-0.391, -0.215]** | **1.000** |
| nsight | -0.010 | 0.028 | [-0.065, 0.046] | 1.000 |
| **CoreHomeRange** | **0.186** | **0.036** | **[0.116, 0.256]** | **1.001** |
| **Diseased (Yes) × DiseasedCount** | **0.115** | **0.045** | **[0.025, 0.202]** | **1.000** |

Note: Estimates are posterior means, with posterior standard deviations (SD), 95% credible intervals (CrI), and convergence diagnostics (R-hat). Positive coefficients indicate greater nearest-neighbour distances, whereas negative coefficients indicate closer spacing. Bold estimates indicate 95% credible intervals that exclude zero.

Importantly, the interaction between focal disease status and diseased conspecific count was positive (posterior mean = 0.12, 95% CrI [0.02 to 0.20]), indicating that the negative association between social distance and diseased conspecifics count was weaker for diseased individuals than for healthy individuals (Fig. 1). Thus, although higher diseased conspecific counts were generally associated with shorter social distances, diseased individuals were observed at relatively greater distances under conditions characterised by higher numbers of diseased conspecifics. Effects for count predictors are expressed per one standard deviation increase.

**Figure 1.**
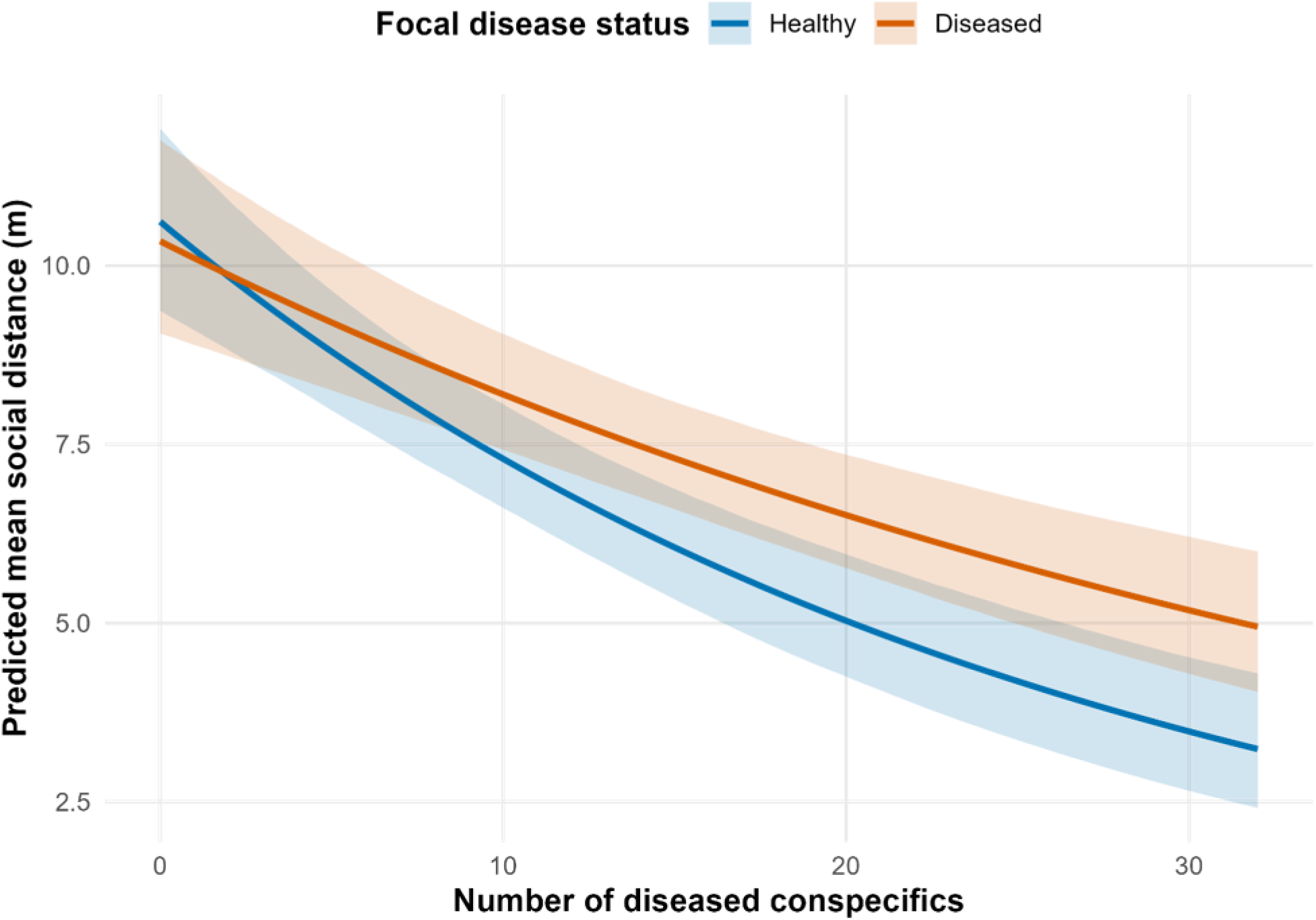
Predicted mean social distance (m) as a function of the number of diseased conspecifics from the Bayesian censored Gamma mixed-effects models. Lines represent posterior mean predictions, and shaded bands indicate 95% credible intervals. Colours denote focal disease status (healthy vs. diseased).

Results were qualitatively consistent under the reduced threshold (≥ 25 sightings), with effect sizes and credible intervals showing no substantive change (Table S4), and the full Bayesian model output (Table S3). Raw observations overlaid on model predictions are provided in Figure S5 to facilitate assessment of the fitted relationship relative to the underlying data.

### Robustness analysis

Additional models were fitted using total conspecific count, healthy conspecifics count, and disease-status categories that distinguished asymptomatic and symptomatic individuals (Table S5-S7). While social distance was negatively associated with total density, consistent with a crowding effect, the interaction with focal disease status was small and not clearly supported, with credible intervals overlapping zero (Table S5). Similarly, when using the number of healthy conspecifics, social distance was negatively associated with healthy conspecific density, but neither the main effect of focal disease status nor the interaction was supported (Table S6). In contrast, a model distinguishing asymptomatic and symptomatic individuals indicated that the interaction with diseased conspecifics count was observed among symptomatic individuals, whereas asymptomatic individuals did not differ from healthy individuals (Table S7).

## Discussion

Our findings indicate that patterns of social distance depend both on an individual’s disease status and on the surrounding disease environment, defined here as the number of diseased conspecifics overlapping the focal individual’s 50% core home range. The model revealed a strong crowding effect, with social distance negatively associated with increasing numbers of diseased conspecifics. However, in areas where more diseased conspecifics overlapped the focal individual’s core home range, the crowding-driven reduction in spacing became weaker. This pattern reflects the fact that diseased individuals were observed at larger distances than healthy individuals under high-disease conditions.

The observed spacing adjustments are consistent with risk-sensitive responses to *Nb*, a pathogen whose environmental persistence enables indirect transmission (Becker et al., 2025). Where pathogens persist in the environment, increased spacing may lower exposure not only during direct encounters but also via shared substrates and space use (Bahram and Netherway, 2022; Becker et al., 2025). Localised clusters of diseased individuals may arise from habitat use or shared microclimatic refugia (Allender et al., 2015; Langwig et al., 2012). Our interaction effect demonstrates that spacing behaviour is shaped jointly by crowding and disease context, rather than by density alone. Comparable clustering and context-dependent spacing have been reported in other environmentally persistent disease systems, such as white-nose syndrome in bats (Langwig et al., 2012), and ophidiomycosis in snakes (Allender et al., 2015).

Additional analysis helps clarify potential explanations for this pattern. Models based on total (Table S5) and healthy conspecific (Table S6) counts revealed strong density-dependent reductions in social distance but did not clearly support disease-dependent interactions, suggesting that general crowding alone does not explain the observed differences. In contrast, the model distinguishing asymptomatic and symptomatic individuals (Table S7) indicated that only symptomatic individuals exhibited distinct spacing patterns in relation to diseased conspecifics, whereas asymptomatic individuals behaved similarly to healthy individuals. Asymptomatic individuals may represent early-stage infection or low infection intensities without visible clinical signs (Grabow et al., 2024), potentially contributing to spacing patterns similar to those of healthy individuals. Although spatial aggregation of diseased individuals was not explicitly quantified, robustness analyses using total and healthy conspecific counts suggest that the observed interaction cannot be explained solely by local density effects.

Shifts in social spacing may arise from multiple, non-exclusive behavioural processes. Increased distances could reflect behavioural changes in diseased individuals themselves, such as reduced social tolerance, active social withdrawal from conspecifics, or movement limitations caused by infection (Lopes et al., 2022). Alternatively, or in combination, they may result from avoidance behaviour by healthy individuals in response to visible signs of disease, consistent with documented disease avoidance strategies across taxa (Behringer et al., 2006; Bos et al., 2012; Heinze and Walter, 2010; Kiesecker et al., 1999; Poirotte et al., 2017; Rueppell et al., 2010). In eastern water dragons, evidence suggests that individuals avoid severely diseased conspecifics (Tacey et al., 2023), indicating that disease-mediated avoidance may contribute to the patterns observed here. However, because our analysis quantifies spatial distance rather than individual-initiated movements, we cannot directly determine whether increased distances arise from healthy individuals avoiding diseased conspecifics, diseased individuals reducing social interactions, or a combination of both mechanisms. Our findings extend these patterns to a free-ranging reptile, demonstrating increased spacing and reduced opportunities for close proximity under elevated disease risk. These results align with the broader predictions that animals modulate social behaviour according to infection risk and environmental cues rather than expressing uniform avoidance (Curtis, 2014).

By analysing population-level patterns in social distance across multiple seasons, we captured longer-term behavioural trends that reflect sustained responses to the combined social and epidemiological environment. This broader temporal scale integrates variation across individuals and time, allowing cumulative, context-dependent effects to emerge, including the attenuation of crowding-driven contraction of spacing and reduced close contact in dense, high-disease settings, patterns that may be difficult to detect in short-term or dyadic studies (Tacey et al., 2023). Although observational and therefore correlational, these findings clarify how disease risk modulates spatial behaviour under natural conditions and complement fine-scale interaction studies. Within a “landscape-of-disgust” framework (Weinstein et al., 2018), our results suggest that disease risk could modify density-driven social behaviour, resulting in nuanced avoidance responses rather than complete social withdrawal.

As the prevalence of an infectious disease rises, individuals may have fewer opportunities to maintain effective social spacing simply because diseased conspecifics become more abundant in the population. This raises the question of whether social spacing remains a viable strategy under high infection pressure, and how these constraints influence transmission and disease dynamics. At the study site, *Nb* prevalence increased from 21.1% in 2018 to 62.4% in 2023; under such conditions, individuals may become progressively less able to maintain effective spacing from diseased conspecifics. This also prompts consideration of the limits of behavioural avoidance in crowded environments. Although season was treated as a random effect and temporal trends were not the primary focus of the present study, the substantial increase in disease prevalence through time suggests that opportunities for social spacing may become increasingly constrained, with potential consequences for transmission dynamics.

Future work should assess whether social strategies remain plastic under escalating pathogen infection pressure and evaluate the consequences for pathogen transmission and host demography. In parallel, understanding how *Nb* persists and accumulates in the environment would provide important insights into transmission risk. Quantifying pathogen survival time and environmental load across habitat types and areas of varying disease prevalence would help determine whether increased behavioural spacing effectively reduces exposure. Such insights are particularly relevant for conservation in urban and fragmented ecosystems, where spatial constraints, environmental reservoirs, and disease risks intersect.

## Acknowledgements

We acknowledge the Turrbal and Yugara people as the First Nations custodians of the lands on which this study was conducted. We pay our respects to their Elders, and to their lores, customs, and creation spirits. We thank the staff at Roma Street Parklands for their ongoing support and cooperation with data collection. We also thank past and present members of the Eastern Water Dragon Project in the Celine Frere Research Group for their assistance with data collection and curation.

## Notes

### Competing Interest Statement

The authors have declared no competing interest.

### Summary of Updates

This version has been revised following peer review. Updates include additional robustness analyses, an exploratory comparison between asymptomatic and symptomatic individuals, revisions to the Methods and Results, expanded supplementary material, and a substantially revised Discussion that addresses alternative explanations and study limitations.

### Funding

- Australian Research Council: FT200100192

