## Supplementary material for "Social Distance and Fungal Disease in an Australian Wild Lizard Population"

**Figure S1**: Representative images of *Nannizziopsis barbatae* infection in eastern water dragons

**Table S1**: Root mean square error (RMSE) and bias across candidate censoring constants

**Table S2**: Collinearity diagnostic (variance inflation factors, VIFs) for the primary censored Gamma mixed-effect model

**Figure S2**: Stan trace plots

**Figure S3**: Posterior predictive checks for censoring behaviour

**Figure S4**: PSIS-LOO cross-validation diagnostics

**Figure S5**: Predicted mean social distance (m) as a function of the number of diseased conspecifics with raw observations overlaid

**Table S3**: Full Bayesian model output for the censored Gamma mixed-effect model examining social distance in eastern water dragons.

**Table S4**: Full Bayesian model output for the censored Gamma mixed-effect model using a minimum of 25 sightings (sensitivity analysis)

**Table S5**: Full Bayesian model output for the censored Gamma mixed-effect model including total conspecific count (TotalCount)

**Table S6**: Full Bayesian model output for the censored Gamma mixed-effect model including healthy conspecific count (HealthyCount)

**Table S7**: Full Bayesian model output for the censored Gamma mixed-effect model including healthy, asymptomatic, and symptomatic disease-status categories

Figure S1: Representative images of *Nannizziopsis barbatae* infection in eastern water dragons. (A) Multifocal moderate to severe yellowish crusted skin lesions affecting the ventral surface of one individual. (B-D) Mild whitish to yellowish skin lesions. (E-G) Moderate to severe proliferative lesions characterised by yellowish crusting on the skin.

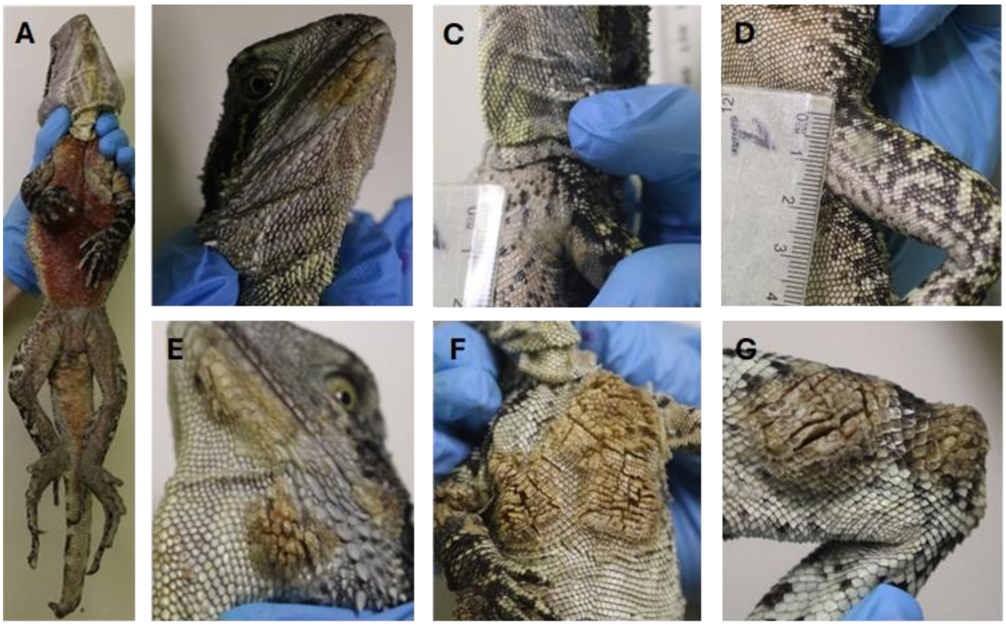

Table S1: RMSE and bias across candidate censoring constants. To ensure numerical stability of the Gamma likelihood while retaining explicit left-censoring, observed values at or below the GPS detection threshold (1.64 m) were replaced with a small positive constant prior to model fitting. The optimal constant was evaluated using simulations based on the fitted intercept-only censored Gamma model (≥30 sightings dataset), with parameter recovery compared across candidate constants ranging from 0.6–1.0 × the detection threshold. Model performance was assessed using RMSE across 30 simulation replicates. The constant equal to the detection threshold (1.0 × 1.64 m) produced the lowest RMSE and was therefore retained for all analyses.

| ratio | mu_bias | mu_rmse |
|---|---|---|
| 0.6 | -0.037100 | 0.087441 |
| 0.7 | -0.037324 | 0.080178 |
| 0.8 | 0.026012 | 0.085895 |
| 0.9 | -0.016704 | 0.077945 |
| **1.0** | **-0.001968** | **0.072352** |

Table S2: Collinearity diagnostic (VIFs) for the primary censored Gamma mixed-effect model. Multicollinearity among fixed effects was assessed using the VIFs, with associated 95% confidence intervals. All predictors exhibited low VIF values, indicating that collinearity was unlikely to influence model inference.

| Parameter | VIF | Increased SE | Tolerance |
|---|---|---|---|
| Sex (Male) | 1.55 | 1.245 | 0.645 |
| Diseased (Yes) | 1.224 | 1.107 | 0.817 |
| DiseasedCount | 2.389 | 1.546 | 0.419 |
| nsight | 1.034 | 1.017 | 0.967 |
| CoreHomeRange | 1.606 | 1.267 | 0.623 |

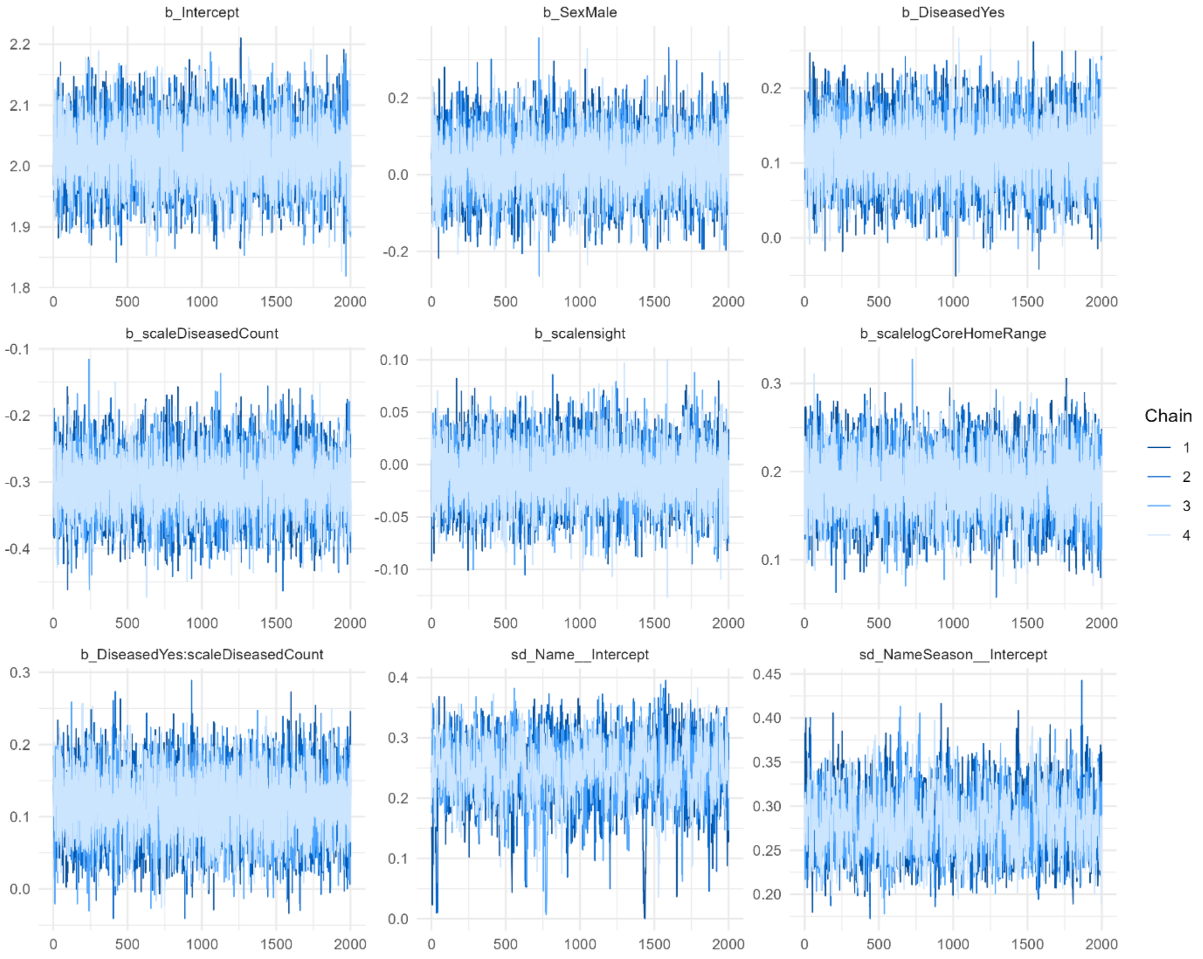


Figure S2: Stan trace plots. Convergence of Markov chain Monte Carlo sampling was assessed using trace plots and the Gelman-Rubin statistic ($\hat{R}$). Visual inspection of trace plots indicated good mixing and stationarity across chains, consistent with satisfactory convergence

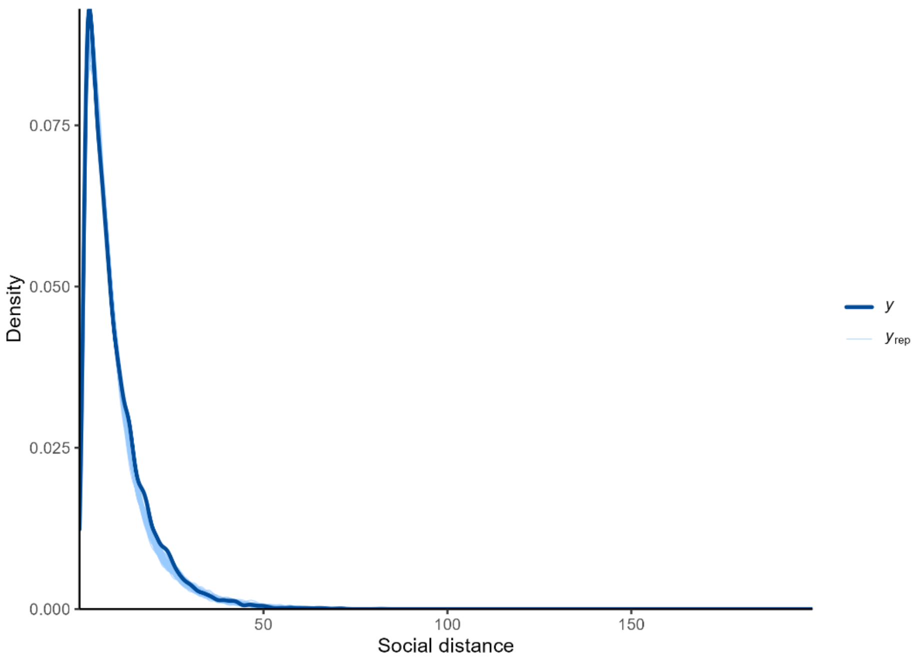


Figure S3: Posterior predictive checks for censoring model. Posterior predictive checks were conducted using replicated data generated from the fitted censored Gamma model. The observed distribution (thick line) is well captured by the posterior predictive distribution (thin lines), including behaviour near the censoring threshold.

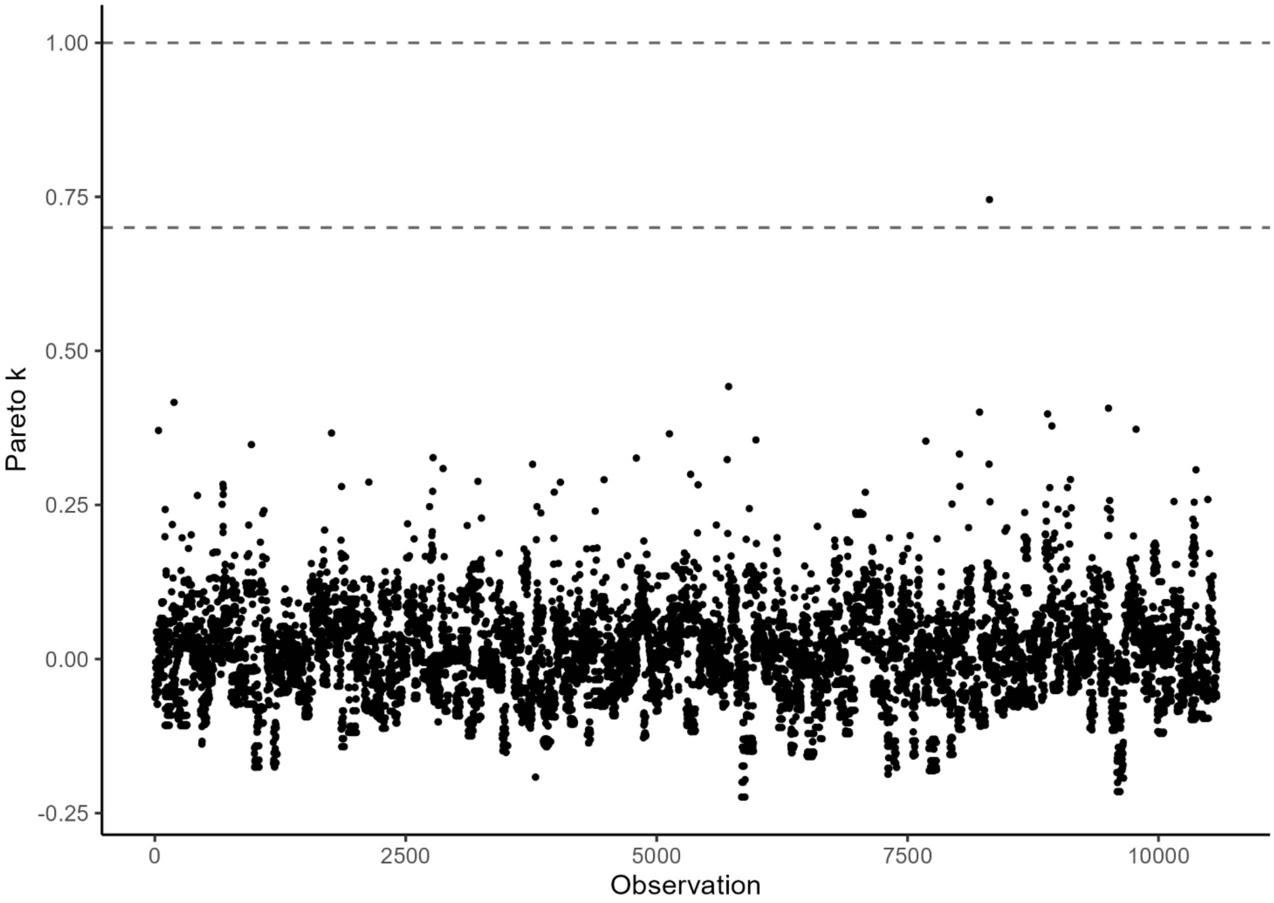


Figure S4: PSIS-LOO cross-validation diagnostics. Pareto-smoothed importance sampling leave-one-out (PSIS-LOO) diagnostics indicated that leave-one-out estimates were stable and reliable.

Figure S5: Predicted mean social distance (m) as a function of the number of diseased conspecifics. Lines represent posterior mean predictions from the Bayesian censored Gamma mixed-effects model and shaded bands indicate 95% credible intervals. Semi-transparent points represent the raw observations used in the analysis. Colours indicate focal disease status (healthy and diseased). For visualisation purposes, the y-axis is restricted to 0-20 m; observations exceeding this range were rare and are omitted from display but retained in the underlying dataset and analysis.

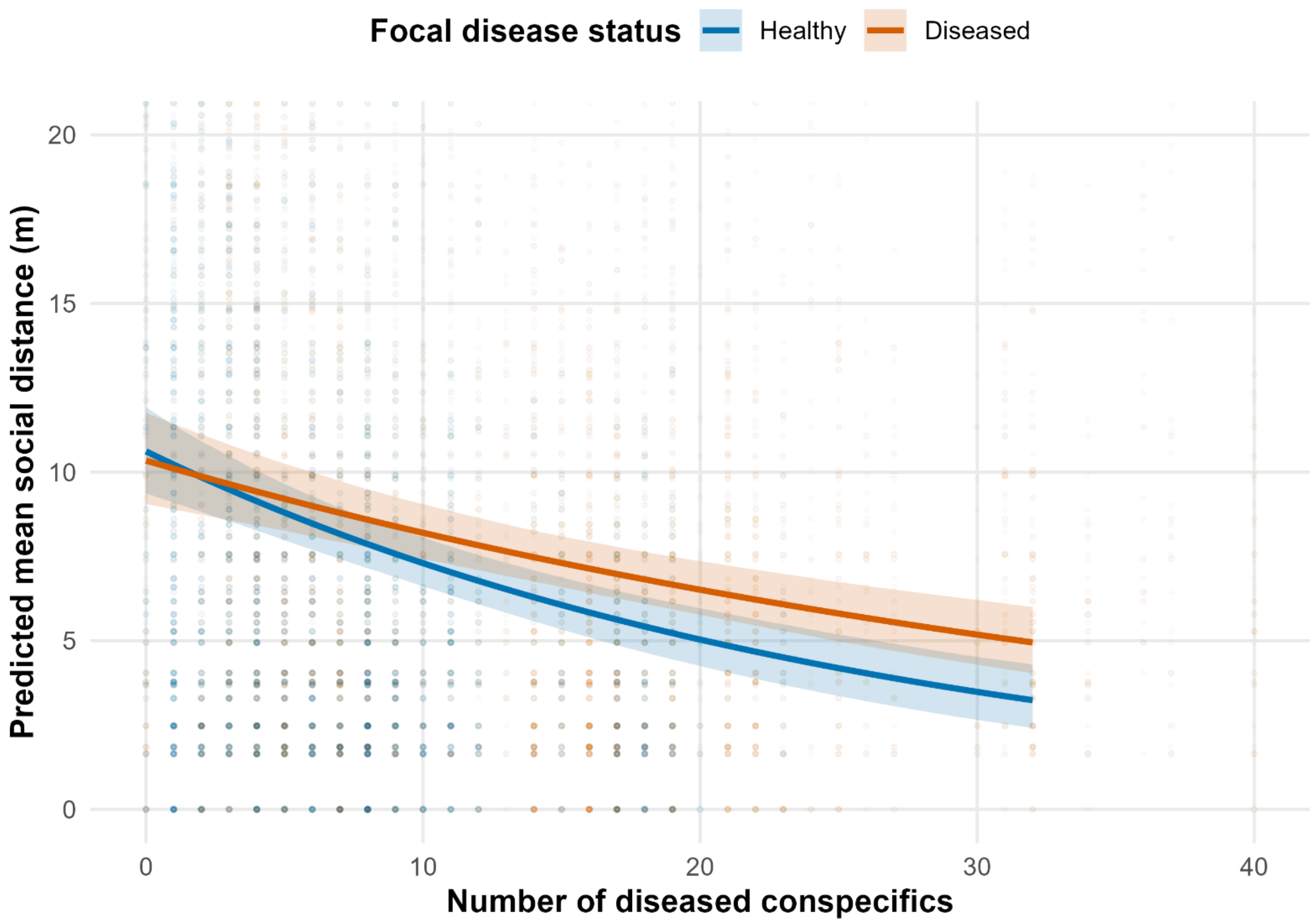

Table S3: Full Bayesian model output for the censored Gamma mixed-effect model examining social distance in eastern water dragons. Posterior summaries are provided for all fixed effects, random-effect variance components, and distributional parameters. Estimates (Posterior mean) are reported with posterior standard errors (Posterior SD), 95% credible intervals (Lower 95% CrI and Upper 95% CrI), convergence diagnostics (R-hat), and effective sample sizes for the bulk and tails of the posterior distribution (Bulk ESS and Tail ESS). This table is provided to facilitate full evaluation of model estimates, uncertainty, and model performance.

| Parameter | Posterior mean | Posterior SD | Lower 95% CrI | Upper 95% CrI | R-hat | Bulk ESS | Tail ESS |
|---|---|---|---|---|---|---|---|
| Intercept | 2.021 | 0.052 | 1.919 | 2.125 | 1.001 | 12681 | 5650 |
| Sex (Male) | 0.031 | 0.076 | -0.118 | 0.18 | 1.001 | 11425 | 6064 |
| Diseased (Yes) | 0.114 | 0.042 | 0.032 | 0.197 | 1.001 | 14498 | 6358 |
| DiseasedCount | -0.303 | 0.045 | -0.39 | -0.213 | 1 | 9597 | 5904 |
| nsight | -0.01 | 0.028 | -0.065 | 0.044 | 1.001 | 12083 | 5982 |
| CoreHomeRange | 0.186 | 0.035 | 0.118 | 0.255 | 1.001 | 7768 | 5644 |
| Diseased (Yes) × DiseasedCount | 0.116 | 0.045 | 0.028 | 0.201 | 1 | 11635 | 6439 |
| Individual identity (Name) SD | 0.246 | 0.05 | 0.135 | 0.331 | 1.001 | 1126 | 1171 |
| Individual-season (Name/Season) SD | 0.281 | 0.033 | 0.222 | 0.352 | 1 | 1471 | 1797 |
| Gamma shape | 1.727 | 0.023 | 1.682 | 1.773 | 1.001 | 12878 | 5072 |

Table S4: Full Bayesian model output for the censored Gamma mixed-effect model using a minimum of 25 sightings (sensitivity analysis). To assess the robustness of model inference to the minimum sighting threshold, all analyses were repeated using a reduced inclusion criterion of ≥25 sightings per individual-season (primary analysis: ≥30 sightings). Model structure and censoring procedures were identical to those used in the primary analysis. Parameter estimates, credible intervals, and effect directions were qualitatively consistent with those of the primary model, indicating that conclusions were robust to variation in sighting threshold. Posterior summaries are provided for all fixed effects, random-effect variance components, and distributional parameters. Estimates (posterior mean) are reported with posterior standard errors (posterior SD), 95% credible intervals (Lower 95% CrI and Upper 95% CrI), convergence diagnostics (R-hat), and effective sample sizes for the bulk and tails of the posterior distribution (Bulk ESS and Tail ESS). This table is provided to facilitate full evaluation of model estimates, uncertainty, and model performance.

| Parameter | Posterior mean | Posterior SD | Lower 95% CrI | Upper 95% CrI | R-hat | Bulk ESS | Tail ESS |
|---|---|---|---|---|---|---|---|
| Intercept | 2.057 | 0.05 | 1.961 | 2.156 | 1 | 21420 | 17750 |
| Sex (Male) | 0.005 | 0.072 | -0.136 | 0.148 | 1 | 21265 | 17154 |
| Diseased (Yes) | 0.116 | 0.039 | 0.04 | 0.192 | 1 | 30213 | 19229 |
| DiseasedCount | -0.294 | 0.041 | -0.374 | -0.211 | 1 | 16940 | 16285 |
| nsight | -0.025 | 0.027 | -0.078 | 0.028 | 1 | 21592 | 17355 |
| CoreHomeRange | 0.201 | 0.034 | 0.133 | 0.267 | 1 | 17299 | 16919 |
| Diseased (Yes) × DiseasedCount | 0.091 | 0.042 | 0.008 | 0.173 | 1 | 23568 | 17538 |
| Individual identity (Name) SD | 0.247 | 0.047 | 0.139 | 0.327 | 1 | 2891 | 2167 |
| Individual-season (Name/Season) SD | 0.278 | 0.032 | 0.221 | 0.346 | 1 | 3405 | 3068 |
| Gamma shape | 1.726 | 0.022 | 1.683 | 1.771 | 1 | 44645 | 16293 |

Table S5: Full Bayesian model output for the censored Gamma mixed-effect model including total conspecific count (TotalCount). Posterior summaries are provided for all fixed effects, random-effect variance components, and distributional parameters. Estimates (Posterior mean) are reported with posterior standard errors (Posterior SD), 95% credible intervals (Lower 95% CrI and Upper 95% CrI), convergence diagnostics (R-hat), and effective sample sizes for the bulk and tails of the posterior distribution (Bulk ESS and Tail ESS). This table is provided to facilitate full evaluation of model estimates, uncertainty, and model performance.

| Parameter | Posterior mean | Posterior SD | Lower 95% CrI | Upper 95% CrI | R-hat | Bulk ESS | Tail ESS |
|---|---|---|---|---|---|---|---|
| Intercept | 2.037 | 0.043 | 1.954 | 2.12 | 1 | 19832 | 22446 |
| Sex (Male) | 0.041 | 0.062 | -0.08 | 0.162 | 1 | 20937 | 22814 |
| Diseased (Yes) | 0.077 | 0.037 | 0.005 | 0.147 | 1 | 32167 | 24984 |
| TotalCount | -0.351 | 0.032 | -0.413 | -0.288 | 1 | 16721 | 22933 |
| nsight | -0.029 | 0.024 | -0.077 | 0.019 | 1 | 18821 | 21259 |
| CoreHomeRange | 0.25 | 0.031 | 0.19 | 0.31 | 1 | 16436 | 22316 |
| Diseased (Yes) × TotalCount | 0.07 | 0.036 | -0.001 | 0.14 | 1 | 22231 | 23753 |
| Individual identity (Name) SD | 0.149 | 0.059 | 0.018 | 0.247 | 1.001 | 1171 | 1938 |
| Individual-season (Name/Season) SD | 0.267 | 0.03 | 0.21 | 0.325 | 1.001 | 1780 | 5027 |
| Gamma shape | 1.728 | 0.023 | 1.682 | 1.773 | 1 | 69959 | 21602 |

Table S6: Full Bayesian model output for the censored Gamma mixed-effect model including healthy conspecific count (HealthyCount). Posterior summaries are provided for all fixed effects, random-effect variance components, and distributional parameters. Estimates (Posterior mean) are reported with posterior standard errors (Posterior SD), 95% credible intervals (Lower 95% CrI and Upper 95% CrI), convergence diagnostics (R-hat), and effective sample sizes for the bulk and tails of the posterior distribution (Bulk ESS and Tail ESS). This table is provided to facilitate full evaluation of model estimates, uncertainty, and model performance.

| **Parameter** | **Posterior mean** | **Posterior SD** | **Lower 95% CrI** | **Upper 95% CrI** | **R-hat** | **Bulk ESS** | **Tail ESS** |
|---|---|---|---|---|---|---|---|
| Intercept | 2.093 | 0.05 | 1.995 | 2.193 | 1 | 8718 | 5668 |
| Sex (Male) | 0.077 | 0.077 | -0.072 | 0.228 | 1 | 8017 | 5891 |
| Diseased (Yes) | -0.022 | 0.039 | -0.098 | 0.054 | 1 | 12670 | 5726 |
| HealthyCount | -0.242 | 0.034 | -0.308 | -0.177 | 1 | 8606 | 6379 |
| nsight | -0.037 | 0.028 | -0.091 | 0.017 | 1 | 6831 | 5664 |
| CoreHomeRange | 0.215 | 0.037 | 0.143 | 0.287 | 1.001 | 5625 | 5813 |
| Diseased (Yes) × HealthyCount | 0.001 | 0.038 | -0.075 | 0.077 | 1 | 11611 | 6781 |
| Individual identity (Name) SD | 0.279 | 0.045 | 0.181 | 0.357 | 1.004 | 1465 | 1348 |
| Individual-season (Name/Season) SD | 0.245 | 0.033 | 0.189 | 0.317 | 1.005 | 1387 | 1428 |
| Gamma shape | 1.728 | 0.023 | 1.682 | 1.775 | 1.002 | 14408 | 4890 |

Table S7: Full Bayesian model output for the censored Gamma mixed-effect model including healthy, asymptomatic, and symptomatic disease-status categories. Posterior summaries are provided for all fixed effects, random-effect variance components, and distributional parameters. Estimates (Posterior mean) are reported with posterior standard errors (Posterior SD), 95% credible intervals (Lower 95% CrI and Upper 95% CrI), convergence diagnostics (R-hat), and effective sample sizes for the bulk and tails of the posterior distribution (Bulk ESS and Tail ESS). This table is provided to facilitate full evaluation of model estimates, uncertainty, and model performance.

| Parameter | Posterior mean | Posterior SD | Lower 95% CrI | Upper 95% CrI | R-hat | Bulk ESS | Tail ESS |
|---|---|---|---|---|---|---|---|
| Intercept | 1.904 | 0.046 | 1.813 | 1.995 | 1 | 22718 | 17772 |
| Sex (Male) | 0.048 | 0.068 | -0.086 | 0.181 | 1 | 20887 | 18152 |
| Diagnostic (Asymptomatic) | -0.006 | 0.063 | -0.13 | 0.118 | 1 | 30629 | 18326 |
| Diagnostic (Symptomatic) | 0.144 | 0.04 | 0.065 | 0.222 | 1 | 30992 | 20330 |
| DiseasedCount | -0.346 | 0.037 | -0.418 | -0.274 | 1 | 17702 | 17153 |
| nsight | -0.032 | 0.026 | -0.082 | 0.018 | 1 | 20905 | 17474 |
| CoreHomeRange | 0.196 | 0.032 | 0.132 | 0.26 | 1 | 18448 | 16180 |
| Diagnostic (Asymptomatic) × DiseasedCount | -0.003 | 0.095 | -0.19 | 0.183 | 1 | 31205 | 19066 |
| Diagnostic (Symptomatic) × DiseasedCount | 0.105 | 0.037 | 0.032 | 0.178 | 1 | 24766 | 18625 |
| Individual identity (Name) SD | 0.217 | 0.049 | 0.102 | 0.299 | 1.001 | 2104 | 1752 |
| Individual-season (Name/Season) SD | 0.284 | 0.03 | 0.23 | 0.346 | 1.001 | 2977 | 2895 |
| Gamma shape | 1.745 | 0.023 | 1.701 | 1.789 | 1 | 46557 | 14571 |
